# α-synuclein preformed fibrils bind to β-neurexins and impair β-neurexin-mediated presynaptic organization

**DOI:** 10.1101/2023.01.28.526024

**Authors:** Benjamin Feller, Aurélie Fallon, Wen Luo, Phuong Trang Nguyen, Irina Shlaifer, Alfred Kihoon Lee, Samer Karkout, Steve Bourgault, Thomas M Durcan, Hideto Takahashi

## Abstract

Synucleinopathies form a group of neurodegenerative diseases defined by misfolding and aggregation of alpha-synuclein (α-syn). Abnormal accumulation and spreading of α-syn aggregates lead to synapse dysfunction and neuronal cell death. Yet, little is known about synaptic mechanisms underlying α-syn pathology. Here we identified β-isoforms of neurexins (β-NRXs) as presynaptic organizing proteins that interact with α-syn preformed fibrils (α-syn PFFs), toxic α-syn aggregates, but not α-syn monomers. Our cell surface protein binding assays and surface plasmon resonance assays reveal that α-syn PFFs bind directly to β-NRX through their N-terminal histidine-rich domain (HRD) at nanomolar range (Kd: ~500 nM monomer equivalent). Furthermore, our artificial synapse formation assays show that α-syn PFFs diminish excitatory and inhibitory presynaptic organization induced by a specific isoform of neuroligin 1 that binds only β-NRXs, but not α-isoforms of neurexins. Thus, our data suggest that α-syn PFFs interact with β-NRXs to inhibit β-NRX-mediated presynaptic organization, providing novel molecular insight into how α-syn PFFs induce synaptic pathology in synucleinopathies such as Parkinson’s disease and dementia with Lewy bodies.

## Introduction

Synucleinopathies form a group of neurodegenerative disorders that includes Parkinson’s disease (PD), PD dementia (PDD), dementia with Lewy bodies (DLB), the Lewy body variant (LBV) of Alzheimer’s disease (AD), and multiple system atrophy (1–3). The main pathohistological feature of synucleinopathies is the neuronal and synaptic accumulation of toxic misfolded and aggregated α-synuclein (α-syn), the principal constituent of protein deposits named Lewy bodies (4). α-syn plays crucial roles in the pathogenesis of synucleinopathies including in neuronal toxicity and synaptic dysfunction (5). In particular, α-syn can be released from one neuron and taken up by other neurons, resulting in spreading of α-syn pathology in the brain (6–9). Evidence regarding trans-synaptic transmission of pathological α-syn (10–14) suggests that synaptic adhesion and/or transmission mechanisms are involved in α-syn pathology but the specific molecular mechanisms of this require further investigation.

Synapse formation, maturation, maintenance, and plasticity are regulated by a series of neuronal adhesion molecules that form “synaptic organizing complexes” (15–20). Synaptic organizing complexes are trans-synaptic adhesion complexes that possess synapse organizing activity by which they can induce pre- and/or post-synaptic differentiation (hereinafter termed “synaptogenic” activity) and thereby act as essential molecular signals for normal synapse function. The neurexin (NRX)-based synaptic organizing complexes have been studied most well (17,21–23), and genetic mutations in NRXs and their binding partners such as neuroligins (NLGs) and leucine-rich repeat transmembrane proteins (LRRTMs) are highly linked with cognitive disorders such as autism and schizophrenia (17,21,24–27). Notably, our recent study has uncovered that NRXs bind directly to amyloid-β (Aβ) oligomers (AβOs), and that this interaction leads to NRX dysregulation and dysfunction (28,29). Importantly, like α-syn, Aβ peptides induce neuronal toxicity, synaptic dysfunction and synaptic loss and exhibit neuron-to-neuron transmission in the brain through a trans-synaptic mechanism (30–37). Furthermore, there is considerable overlap of histopathological features in AD and PDD patients: patients with PDD tend to have a high burden of Aβ plaques (38,39), the hallmark pathology of AD, and up to 50% of AD patients present with α-syn pathology (40–42). The strong clinical association and biological similarities between the α-syn and Aβ pathologies suggest that α-syn and Aβ may share common synaptic mechanisms in the progression of their pathologies, with synaptic organizing complexes, especially NRXs, being strong candidates for molecules underlying α-syn pathology.

In this study, we prepared recombinant α-syn preformed fibrils (α-syn PFFs) and performed a candidate screen to isolate α-syn PFF-binding synaptic organizers. We found that α-syn PFFs bind to β-isoforms of NRXs (β-NRXs) with the N-terminal histidine-rich domain (HRD) of β-NRXs being necessary for the binding. Surprisingly, this is the same domain that is responsible for AβO-β-NRX interaction (28,29). Further, we discovered that α-syn PFFs diminish β-NRX-mediated presynaptic organization. Given the role of β-NRXs in synaptic transmission and plasticity (43), our results indicate that β-NRXs may act as an α-syn PFF receptor to mediate α-syn PFF-induced synaptic pathology and that the HRD of β-NRXs has the unique molecular property of binding the pathological protein aggregates that are central to two major types of neurogenerative disease.

## Results

### Screening synaptic organizing molecules for interaction with α-synuclein preformed fibrils isolates neurexin 1β as a candidate α-synuclein binding partner

To test whether and which synaptic organizers bind to α-syn PFFs, we first generated recombinant human α-synuclein and prepared preformed fibrils (PFFs) (**Fig. 1**). By electron microscopy (**Fig. 1A**) and dynamic light scattering analysis (**Fig. 1B**), we confirmed that the approximate size of α-syn PFFs is around 50-60 nm, similar to the size of α-syn PFFs used in previous studies (44–46). By using a Thioflavin-T assay, we further confirmed that α-syn PFFs contained amyloid β-sheet structures (**Fig. 1C**) As reported before (46), the treatment of cultured hippocampal neurons with the prepared α-syn PFFs, but not α-syn monomers, caused a significant increase in the phosphorylation of α-syn in neurites (**Fig. 1D**), indicating that the prepared α-syn PFFs have toxic properties, as expected. To perform cell surface protein binding assays with a good signal-to-noise ratio (**Fig. 2**), we conjugated biotin molecules to the α-syn PFFs to allow us to specifically label bound α-syn PFFs using fluorophore-conjugated streptavidin. We first confirmed the biotin conjugation to α-syn PFFs and α-syn monomers by dot blot analysis (**Fig. 2A**). We further confirmed that the biotin-conjugated α-syn PFFs (biotin-α-syn PFFs) could bind to COS-7 cells expressing the cellular prion protein (PrP^c^), a known α-syn PFF-interacting membrane protein (47–49), but not to those expressing CD4, a negative control protein (**Fig. 2B**). In cell surface protein binding assays using the biotin-α-syn PFFs, we screened a total of 22 synaptic organizers and detected significant binding of biotin-α-syn PFFs to COS cells expressing neurexin1β (NRX1β), but not to those expressing any of the other tested synaptic organizers (**Fig. 2C**). Thus, we isolated NRX1β as a candidate synaptic organizer that binds to α-syn PFFs.

**Figure 1.**
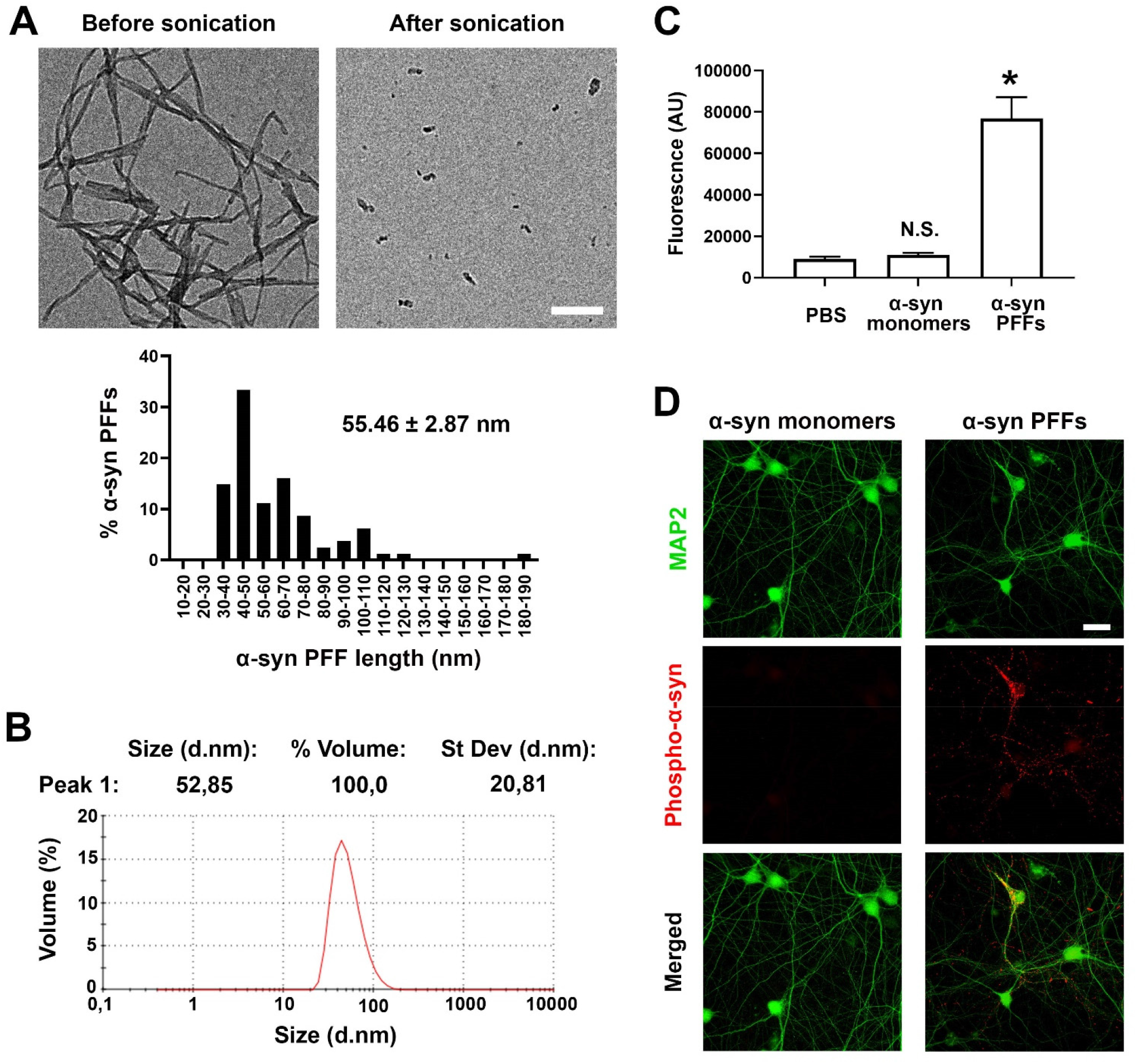
Biophysical characterization of pathological α-synuclein preformed fibrils (α-syn PFFs). (**A**) Representative electron microscopy (EM) images (top) of α-synuclein (α-syn) proteins after aggregation incubation (left) and then sonication (right). The sonicated aggregates are what was used as α-syn preformed fibrils (α-syn PFFs) in this study. Histogram (bottom) showing the lengths of α-syn PFFs. Most are between 30 and 80 nm (55.46 ± 2.87 nm (mean ± SEM, 81 α-syn PFF particles)). (**B**) Dynamic light scattering analysis of α-syn PFFs confirms that their average size is 52.85 nm, consistent with the above EM analysis. (**C**) Fluorescent quantification using thioflavin-T assays to confirm amyloid β-sheet structures in α-syn PFFs. Kruskal-Wallis one-way ANOVA, P < 0.01. *P < 0.05 and N.S., not significant compared with PBS by Dunn’s multiple comparisons test. Data are presented as mean ± SEM. (n = 2 samples for PBS and α-syn monomers and 6 samples for α-syn PFFs). (**D**) α-syn PFFs induce hyperphosphorylation of endogenous α-synuclein in the hippocampal neuron cultures. Hippocampal cultured neurons were treated with α-syn PFFs (2.5 μg/ml [174 nM monomer equivalent]) or α-syn monomers (2.5 μg/ml) at 7 days *in vitro* (DIV). The cultures were maintained until 14 DIV and then immunostained for phospho-α-synuclein (red) and MAP2 (green), the dendrite marker. Treatment with α-syn PFFs, but not α-syn monomers, induced clusters of phosphorylated α-synuclein immunoreactivity along neurites (middle panel, red). Scale bars: 200 nm (**A**) and 30 μm (**C**).

**Figure 2.**
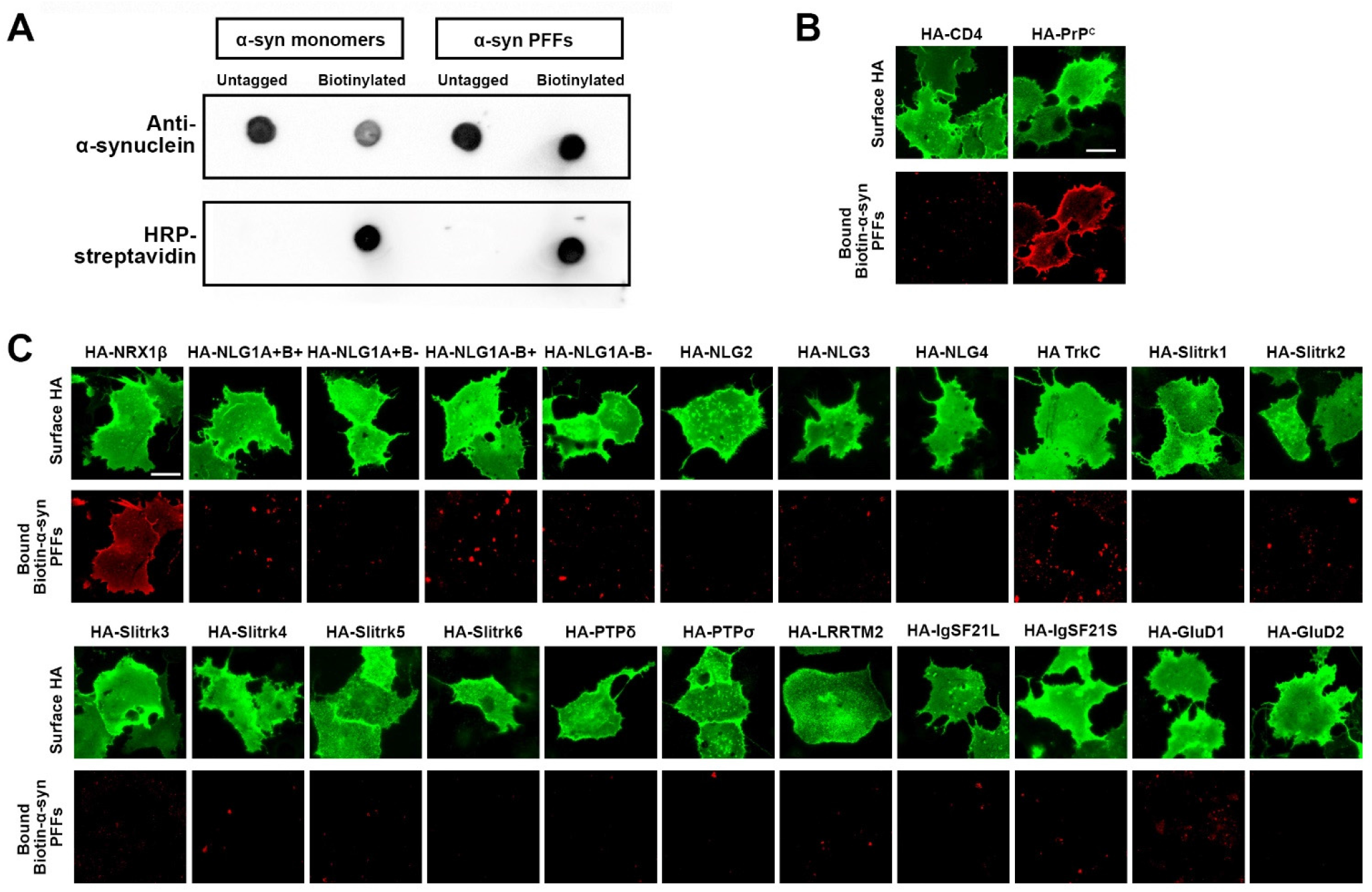
A candidate screen based on cell surface protein binding assays using biotin-conjugated α-syn PFFs isolates neurexin1β as an α-syn PFF-interacting protein. (**A**) Validation of biotin conjugation to α-syn monomers and PFFs by dot blot assays. Untagged and biotin-conjugated α-syn monomers and PFFs were spotted onto a nitrocellulose membrane and immunolabeled with anti-α-synuclein antibody to confirm the presence of the indicated proteins. After stripping the anti-α-synuclein antibody, the membrane was labelled with HRP-conjugated streptavidin to detect biotin-conjugated proteins. Only biotin-α-syn monomers and PFFs, but not untagged ones, display HRP-streptavidin-based signals. (**B**) A cell surface protein binding assay shows the biotin-α-syn PFFs bind to COS-7 cells expressing the N-terminal extracellular HA-tagged cellular prion protein (HA-PrP^c^), which is a known α-syn PFF-interreacting protein, but not those expressing HA-CD4 as a negative control. Surface HA was immunolabelled to verify expression of these constructs on the COS-7 cell surface. (**C**) Representative images showing cell surface protein binding assays testing for interaction between α-syn PFFs (1 μM, monomer equivalent) and known synaptic organizers. COS-7 cells expressing the indicated construct were exposed to α-syn PFFs. α-syn PFFs bind to COS-7 cells expressing HA-neurexin (NRX)1β, but not to those expressing any of the other organizers including HA-neuroligin1 (HA-NLG1). Surface HA (green) was immunostained to verify expression of the indicated N-terminal extracellular HA-tagged constructs on the COS-7 cell surface. Scale bars: 30 μm (**B, C**).

### α-synuclein PFFs, but not α-synuclein monomers, bind directly to neurexin 1β

In cell surface protein binding assays, we next perform saturation analysis and Scatchard plot analysis to test whether the binding between α-syn PFFs and NRX1 β had properties typical of ligand-receptor binding and to determine the binding affinity, respectively (**Fig. 3**). The binding of biotin-α-syn PFFs to COS-7 cells expressing NRX1β showed an increasing and saturable binding curve with increasing amounts of biotin-α-syn PFFs (**Fig. 3A and B**). The dissociation constant (K_D_) value was 536 nM monomer equivalent (**Fig. 3C**). In the cell surface protein binding assays, it remains possible that some endogenous proteins expressed in COS-7 cells might affect and/or mediate an interaction between α-syn PFFs and NRX1β. To test whether they have a direct protein-protein interaction, we further performed surface plasmon resonance (SPR) assays using untagged purified α-syn PFFs and purified recombinant NRX1β ectodomain fused to the human immunoglobin Fc region (NRX1β-Fc) immobilized on the surface of a sensor chip (**Fig. 3D and E**). Association and dissociation rate constants were determined using a 1:1 Langmuir binding model. The SPR sensorgrams of different concentrations of α-syn PFFs showed significant binding responses upon increasing α-syn PFF concentration (**Fig. 3D**). The K_D_ value of α-syn PFF-NRX1β binding measured by the SPR assays was 554 nM monomer equivalent (**Fig. 3D**), consistent with the K_D_ value measured by Scatchard plot analysis in the cell surface protein binding assays (**Fig. 3C**). In contrast, untagged purified α-syn monomers failed to show typical binding responses in the SPR analysis (**Fig. 3E**), indicating negligible binding between α-syn monomers and NRX1β. Together, these results suggest that α-syn PFFs, but not α-syn monomers, bind directly to NRX1β in the nanomolar range under both cell-free and cell-based conditions.

**Figure 3.**
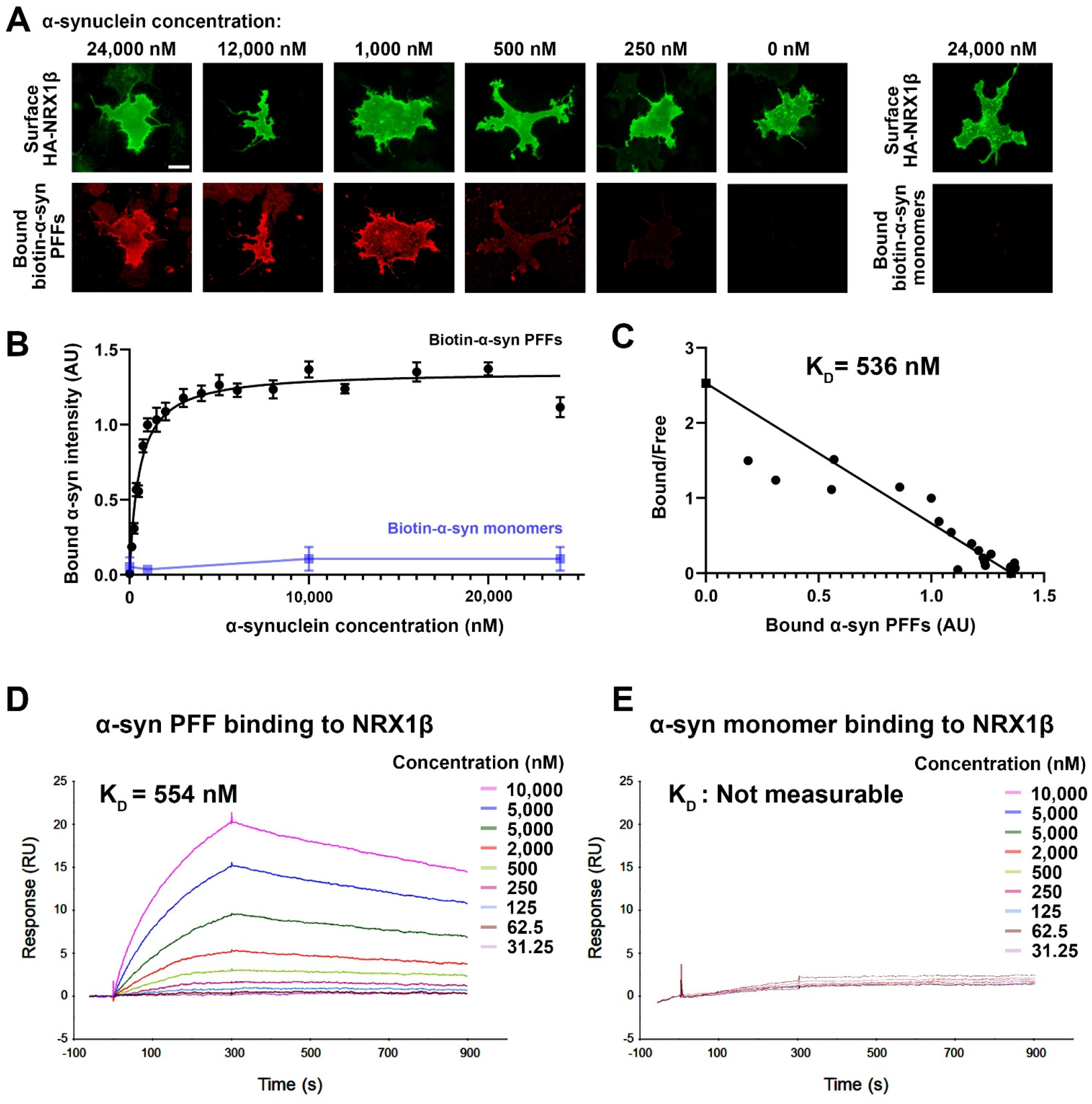
α-syn PFFs, but not α-syn monomers, bind directly to neurexin1β. (**A**) Representative images of cell surface binding assays showing COS-7 cells expressing extracellular HA-tagged neurexin 1 β (HA-NRX1 β) treated with biotin-conjugated α-syn PFFs at the indicated concentrations (0 - 24,000 nM monomer equivalent) or treated with 24,000 nM biotin-α-syn monomers. Surface HA (green) was immunolabelled to verify expression of these constructs on the COS-7 cell surface. Scale bar: 30 μm. (**B**) Saturable binding of biotin-α-syn PFFs to COS-7 cells expressing HA-NRX1β. Biotin-α-syn PFFs display a saturable binding curve whereas biotin-α-syn monomers show no binding even at high concentrations. Data are presented as mean ± SEM (n = 30 cells for each plot from three independent experiments). (**C**) A Scatchard plot of the biotin-α-syn PFF binding data from (**B**) indicates a K_D_ of 536 nM monomer equivalent. (**D, E**) Representative sensorgrams from surface plasmon resonance (SPR) analysis for the binding of untagged α-syn PFFs (**D**) or untagged α-syn monomers (**E**) to purified recombinant NRX1β ectodomain fused to human immunoglobulin Fc (NRX1β-Fc) immobilized on the sensor chip. The K_D_ value of α-syn PFFs is 554 nM. The K_D_ value of α-syn monomers could not be determined, and the sensorgrams indicated no significant binding of α-syn monomers to immobilized NRX1β.

### α-synuclein PFFs specifically bind to β-isoforms of neurexin

NRXs are encoded by three different genes (*NRXN1*, 2 and 3), and each gene possesses two independent promotors that drive the expression of longer α-isoforms (α-NRXs) and shorter β-isoforms (β-NRXs) (17,21,50). α/β-NRXs contain multiple alternative splicing sites, and the splicing site 4 (S4) common to α/β-NRXs is crucial for the selectivity and specificity of NRX-interacting proteins such as NLGs and LRRTMs (17,21,22,50). Therefore, we next tested which NRX isoforms can bind to α-syn PFFs by using cell surface protein binding assays (**Fig. 4**). Biotin-α-syn PFFs strongly bound to NRX1β and NRX2β and weakly bound to NRX3β regardless of S4 insertion (**Fig. 4A and B**). Biotin-α-syn PFFs did not bind to any isoforms of α-NRXs (**Fig. 4A and B**). These results indicate that α-syn PFFs specifically bind to β-isoforms of NRXs.

**Figure 4.**
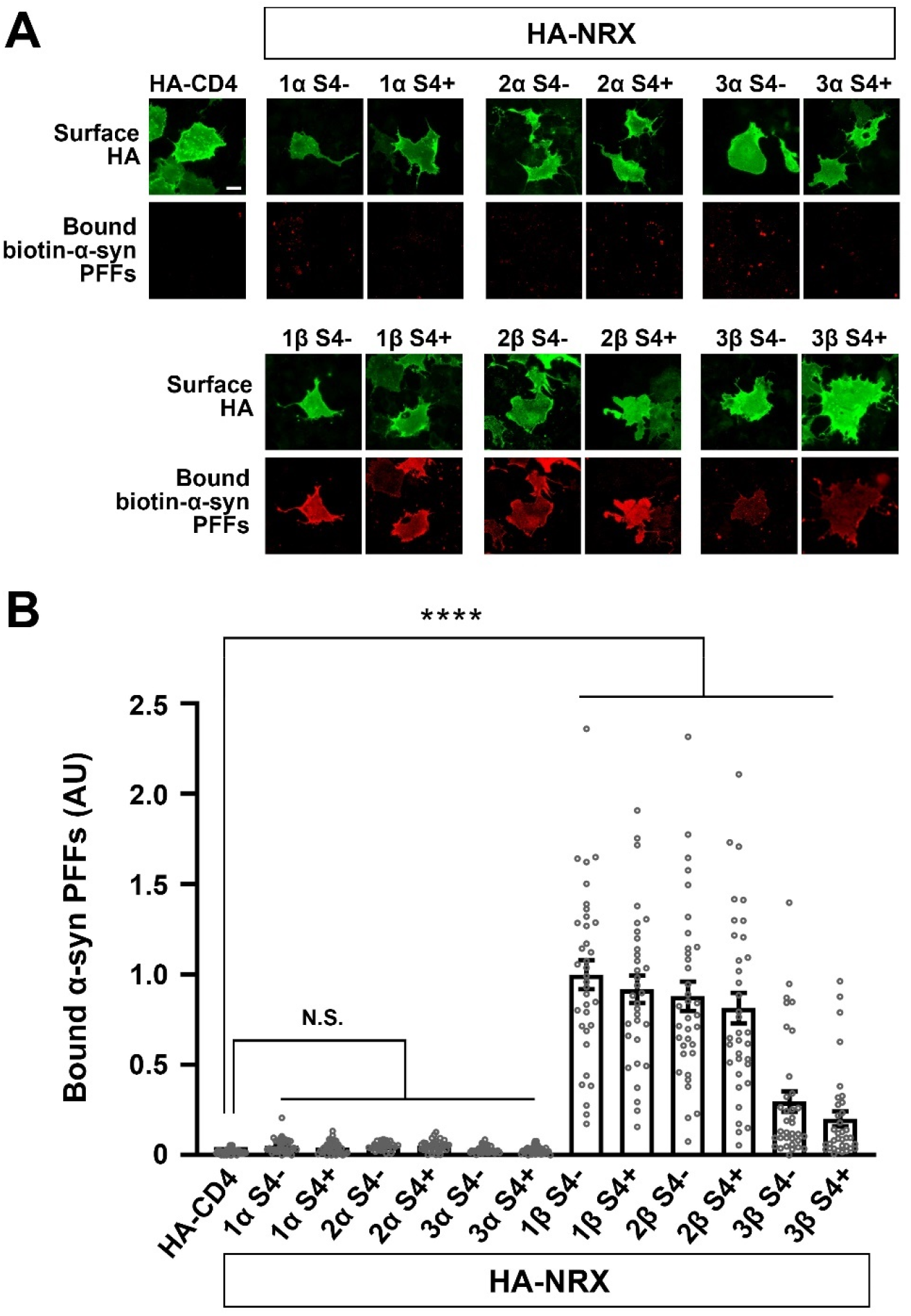
α-syn PFFs selectively bind to β-isoforms, and not α-isoforms, of neurexins. (**A**) Representative images showing the binding of biotin-α-syn PFFs (1 μM, monomer equivalent) to COS-7 cells expressing the indicated isoform of extracellularly HA-tagged NRX. S4- and S4+ indicate without and with an insert at splicing site 4, respectively. Surface HA (green) was immunolabelled to verify expression of these constructs on the COS-7 cell surface. Scale bar: 30 μm. (**B**) Quantification of the average intensity of bound biotin-α-syn PFFs on COS-7 cells expressing the indicated constructs. Only β-isoforms of NRXs show significant binding of biotin-α-syn PFFs. α-NRXs have no binding of biotin-α-syn PFFs. Kruskal-Wallis one-way ANOVA, P < 0.0001. ****P < 0.0001 compared with HA-CD4 by Dunn’s multiple comparisons test. N.S., not significant. Data are presented as mean ± SEM. (n = 30 cells for each from three independent experiments).

### The N-terminal histidine-rich domain of β-neurexins is responsible for the binding of α-synuclein PFFs

β-NRXs possess an N-terminal domain called the histidine-rich domain (HRD) that is absent from α-NRXs (50) (**Fig. 5A**), and this difference could account for the β-NRX binding selectivity of α-syn PFFs. Indeed, we have previously shown that the HRD of β-NRXs is responsible for the binding of AβOs to β-NRXs (28,29). We therefore next tested whether the HRD is responsible for α-syn PFF binding by using NRX1β deletion constructs in cell surface protein binding assays (**Fig. 5**). Deletion of the HRD from NRX1β completely abolished the binding of biotin-α-syn PFFs (**Fig. 5A-C**). However, the binding of biotin-α-syn PFFs was not affected by deletion of the LNS domain, which is responsible for the binding of postsynaptic ligands such as NLGNs and LRRTMs, nor by deletion of the cysteine loop region or by introduction of the point mutation that prevents heparan sulfate modification of NRXs (**Fig. 5A-C**). There was no difference among NRX1β deletion constructs in surface HA expression (**Fig. 5D**), indicating that the lack of binding of biotin-α-syn PFFs to NRX1β ΔHRD was not due to insufficient surface expression of NRX1β ΔHRD. Deletion of the HRD from NRX2β and NRX3β also diminished the binding of biotin-α-syn PFFs to NRX2β and 3β, respectively (**Supplementary Figure 1**). Thus, the domain analysis indicates that the HRD is a responsible domain for α-syn PFF binding to β-NRXs. We further confirmed the involvement of the HRD in α-syn PFF binding using an antibody that recognizes an epitope in the NRX1β HRD. COS-7 cells expressing NRX1β were exposed to a single concentration (1 μM monomer equivalent) of biotin-α-syn PFFs in the presence of varying concentrations (0 – 5 μg/ml) of anti-NRX1β antibody (**Fig. 5E and F**). Treatment with the anti-NRX1β antibody inhibited the binding of biotin-α-syn PFFs to NRX1 β in a dose-dependent manner (**Fig. 5F**). The inhibition curve indicated that the half-maximal inhibition concentration (IC_50_) value for anti-NRX1β antibody was 0.68 μg/ml (**Fig. 5F**). These data also support our conclusion that the HRD is responsible for the binding of α-syn PFFs to β-NRXs.

**Figure 5.**
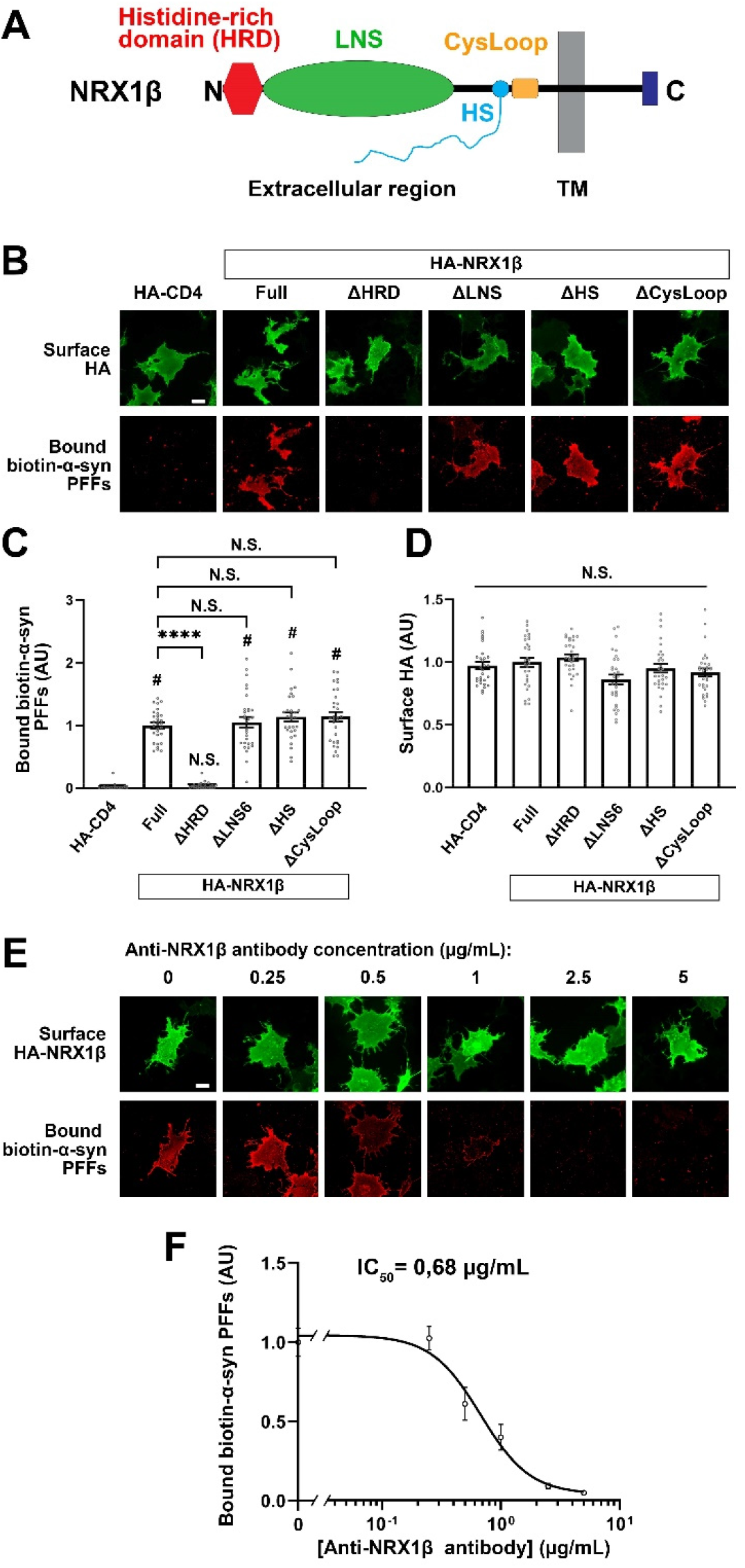
The N-terminal histidine-rich domain of neurexin1β is responsible for the binding of α-syn PFFs to neurexin1β. (**A**) A diagram showing the domain structure of NRX1β. HRD: the histidine-rich domain (HRD), LNS: laminin-neurexin-sex hormone binding globulin, HS: a heparan sulfate modification site, CysLoop: cysteine loop region, TM: transmembrane region, N and C: N- and C-terminals, respectively. (**B**) Representative images showing the binding of 1 μM biotin-α-syn PFFs to COS-7 cells expressing the indicated HA-NRX1β deletion constructs, full-length HA-NRX1β or HA-CD4, a negative control. NRX1β lacking the HRD (ΔHRD) has no binding of biotin-α-syn PFFs. The binding of biotin-α-syn PFFs to NRX1β lacking either the LNS, the HS or the CysLoop (ΔLNS, ΔHS, ΔCysLoop) appears comparable to the binding to full-length NRX1 β. (**C**) Quantification of the average intensity of bound biotin-α-syn PFFs on COS-7 cells expressing the indicated HA-NRX1β constructs. Kruskal-Wallis one-way ANOVA, P < 0.0001. ****P < 0.0001 compared with HA-CD4 by Dunn’s multiple comparisons test. N.S., not significant. Data are presented as mean ± SEM. (n = 30 cells for each from three independent experiments). (**D**) Quantification of the average intensity of surface HA on COS-7 cells expressing the indicated HA-NRX1β constructs in (**C**). Kruskal-Wallis one-way ANOVA, P = 0.0034. N.S., not significant in the comparisons with HA-CD4 and full-length HA-NRX1β. Data are presented as mean ± SEM. (n = 30 cells for each from three independent experiments). (**E**) Representative images of cell surface protein binding assays showing COS-7 cells expressing HA-NRX1β co-treated with 1 μM biotin-α-syn PFFs and a mouse monoclonal antibody against the histidine-rich domain (HRD) of neurexin1β (anti-NRX1β antibody) at varying concentrations (0 – 5 μg/ml). Cotreatment with anti-NRX1β appears to inhibit the binding of α-syn PFFs to COS-7 cells expressing HA-NRX1 β in a dose-dependent manner. (**F**) Quantification of biotin-α-syn PFFs (1 μM monomer equivalent) bound to COS-7 cells expressing HA-NRX1 β in the presence of various concentrations of anti-NRX1 β antibody (0 – 5 μg/ml). The half maximal inhibitory concentration (IC_50_) is 0.68 μg/ml. Scale bars: 30 μm (**B** and **E**).

### Neuronal overexpression of NRX1β enhances the binding of α-syn PFFs on the axon surface in an NRX1β HRD-dependent manner

Given that β-NRXs are predominantly expressed at presynaptic terminals and in axons (51–53), we next tested whether the binding of α-syn PFFs to β-NRXs occurs on the axon surface as it does on the COS-7 cell surface by performing experiments in neurons overexpressing NRX1β (**Fig. 6**). Primary hippocampal neurons were first transfected to express extracellular HA-tagged NRX1β (HA-NRX1β) and GFP (HA-NRX1β-IRES-GFP), HA-NRX1β lacking its HRD and GFP (HA-NRX1βΔHRD-IRES-GFP), or only GFP (IRES-GFP). These cultures were exposed to biotin-α-syn PFFs and then labelled for bound biotin-α-syn PFFs, surface HA, and MAP2, a dendrite marker, to identify the GFP-positive and MAP-negative neurites as axons (**Fig. 6A**). We then measured the signal intensity of biotin-α-syn PFFs bound on the axons (**Fig. 6B**). In neuron cultures transfected with IRES-GFP alone (as a baseline control), a few axons showed punctate signals corresponding to bound biotin-α-syn PFFs, but this was undetectable in most axons (**Fig. 6A and B**). In contrast, almost all axons transfected with HA-NRX1β-IRES-GFP had strong signals for bound biotin-α-syn PFFs (**Fig. 6A and B**). This enhanced axonal binding of biotin-α-syn PFFs was not observed in neurons transfected with HA-NRX1βΔHRD-IRES-GFP (**Fig. 6A and B**), even though surface expression of HA-NRX1βΔHRD on axons was comparable to that in cultures expressing HA-NRX1β (**Fig. 6C**). These results suggest that NRX1β can mediate α-syn PFF binding to the axon surface through its HRD.

**Figure 6.**
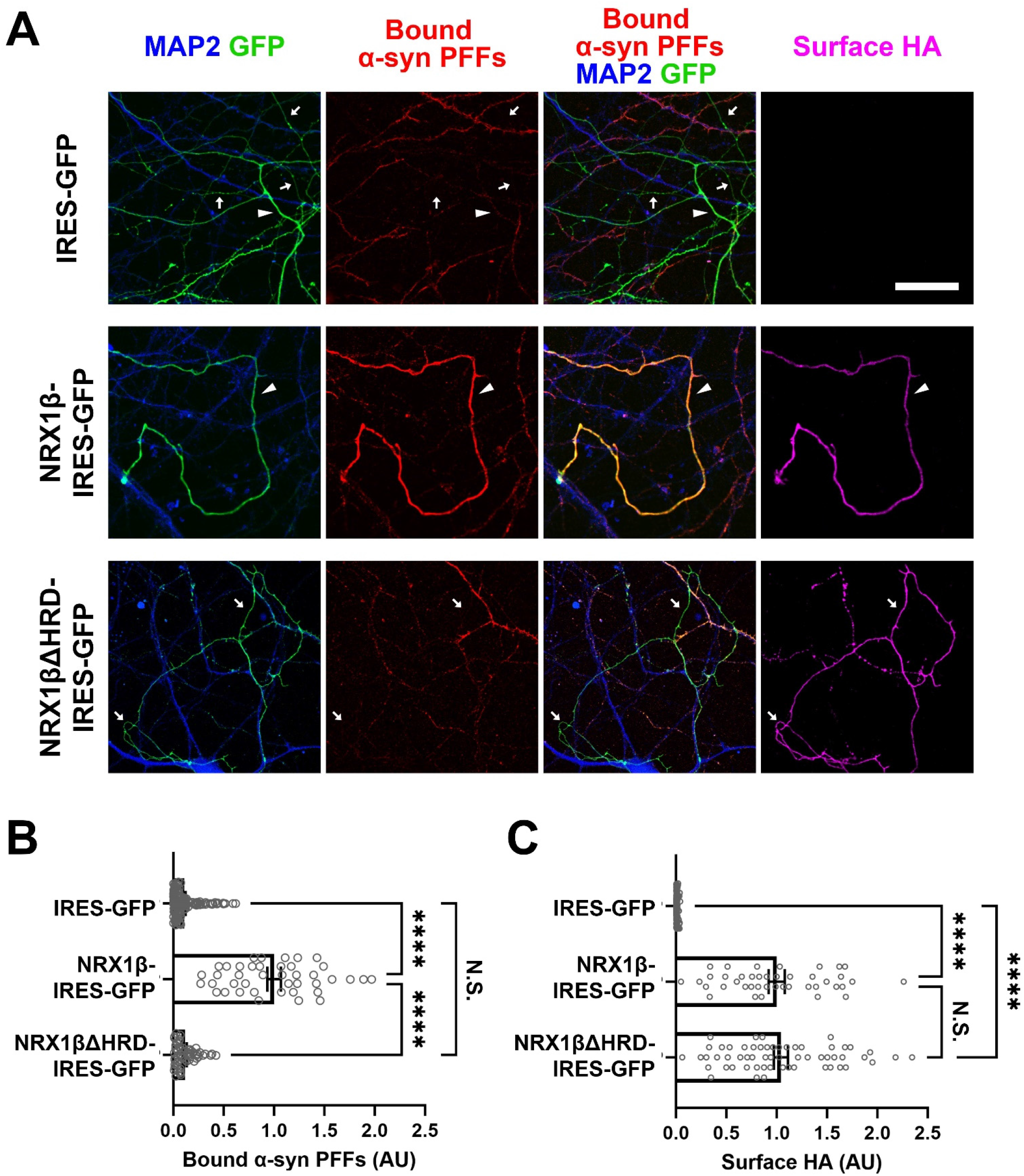
Axonal expression of NRX1β enhances the binding of α-syn PFFs to axon surface in its HRD-dependent manner. (**A**) Representative images of cultured hippocampal neurons transfected with IRES-GFP, HA-NRX1β-IRES-GFP or HA-NRX1βΔHRD-IRES-GFP and exposed to biotin-α-syn PFFs (174 nM monomer equivalent) at 21 DIV. Afterwards, the neurons were triple immunostained for bound biotin-α-syn PFFs, surface HA and MAP2. GFP-positive and MAP2-negative neurites were selected as axons for quantitative analysis. The arrowheads and arrows indicate axons with and without bound α-syn PFF signals, respectively. In the IRES-GFP images (upper), the arrowhead indicates the axon with weak and small punctate bound α-syn PFFs signal. Scale bar: 30 μm. (**B**) Quantification of the average intensity of biotin-α-syn PFFs bound to axons of cultured hippocampal neurons transfected with the indicated constructs. Kruskal-Wallis one-way ANOVA, P < 0.0001. ****P < 0.0001 in the indicated comparisons by Dunn’s multiple comparisons test. N.S., not significant. Data are presented as mean ± SEM. (n = 137, 41 and 58 axons for IRES-GFP, HA-NRX1β-IRES-GFP or HA-NRX1βΔHRD-IRES-GFP, respectively, from three independent experiments). (**C**) Quantification of the surface HA expression in the cultured hippocampal neurons transfected with the indicated constructs and analyzed in (**B**). The surface HA expression level in the analyzed axons was comparable between HA-NRX1β and HA-NRX1βΔHRD, suggesting that the lack of significant binding of biotin-α-syn PFFs onto axons transfected with NRX1βΔHRD-IRES-GFP is not due to insufficient surface expression of NRX1βΔHRD but rather due to the lack of the HRD, which responsible for α-syn PFF binding. Kruskal-Wallis one-way ANOVA, P < 0.0001. ****P < 0.0001 in the indicated comparisons by Dunn’s multiple comparisons test. N.S., not significant.

### α-syn PFF treatment diminishes β-NRX-mediated presynaptic organization

One of the key functions of NRXs expressed on axons is to mediate presynaptic organization trans-synaptically induced by NRX-binding partners such as NLGs (15–18,21,22,54). Therefore, we next tested whether and how α-syn PFFs affect NRX-mediated presynaptic organization by performing artificial synapse formation assays using co-cultures of HEK293T cells expressing NLG1 or NLG2 and primary hippocampal neurons (**Fig. 7**). We chose the isoforms of NLG1 and NLG2 based on the functions of the splicing inserts. The insertion at splicing site B in NLGs prevents their binding to α-NRXs whereas NLG splicing site A does not regulate NRX binding (22,55). We used a NLG1 isoform lacking the splicing site A insert but possessing the splicing site B insert (NLG1A-B+), which binds to only β-NRXs and not α-NRXs (**Supplementary Figure 2A**), and NLG2A+, a NLG2 isoform possessing the splicing site A insert (but which always lacks the splicing site B insert) that binds to both α- and β-NRXs (**Supplementary Figure 2B**) and is endogenously more abundant than the NLG2 isoform lacking the splicing site A (56). In vehicle-treated co-cultured hippocampal neurons, HEK cells expressing NLG1A-B+ or NLG2A+ induced significant accumulation of the excitatory presynaptic marker VGLUT1 (**Fig. 7A-C**) and of the inhibitory presynaptic marker VGAT (**Fig. 7D-F**). Treatment with α-syn PFFs almost fully blocked the NLG1A-B+-induced VGLUT1 (**Fig. 7A-C**) and VGAT accumulation (**Fig. 7D-F**) without altering surface expression of HA-NLG1A-B+ on the HEK cells (**Supplementary Figure 3**). In contrast, α-syn PFF treatment had no effect on NLG2A+-induced accumulation of VGLUT1 (**Fig. 7A-C)** or VGAT (**Fig. 7D-F**), and treatment with α-syn monomers had no effect on VGLUT1 or VGAT accumulation induced by either NLG1A-B+ or NLG2A+ (**Fig. 7**). These results suggest that α-syn PFFs selectively diminish β-NRX-mediated presynaptic organization, which is consistent with our cell surface protein binding assays showing selective binding of α-syn PFFs to β-NRXs but not α-NRXs.

**Figure 7.**
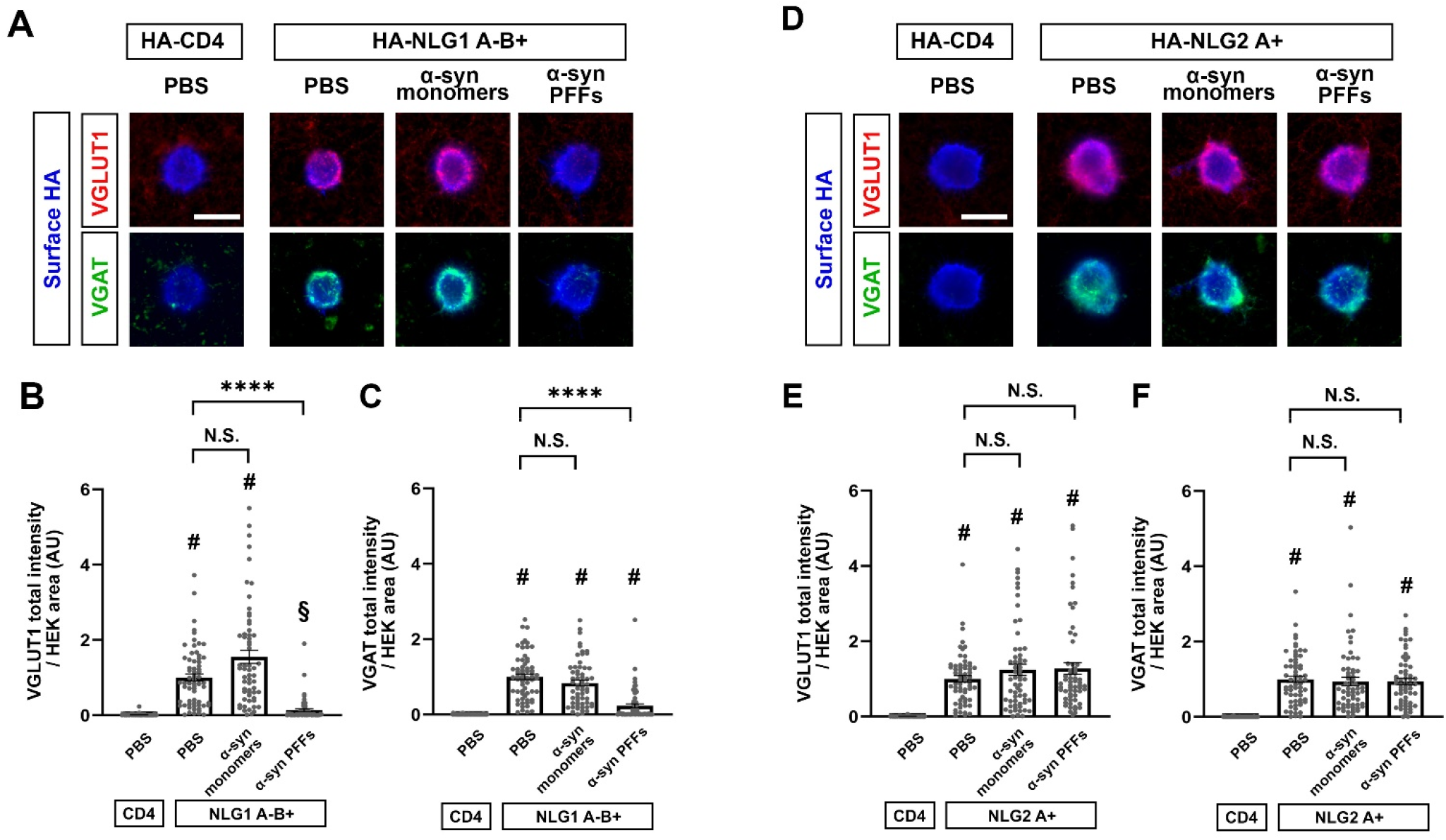
Co-culture-based artificial synapse formation assays reveal that α-syn PFFs diminish presynaptic differentiation induced by the NLG1 A-B+ isoform. (**A, D**) Representative images from an artificial synapse formation assay showing the accumulation of VGLUT1, an excitatory presynaptic vesicle protein, and VGAT, an inhibitory presynaptic vesicle protein, induced by HA-NLGN1 that lacks the splicing site A insert but possesses one at splicing site B (HA-NLGN1A-B+; **A**) or by HA-NLG2 that possesses the splicing site A insert (HA-NLG2A+; **D**). HA-CD4 was used as a negative control. α-syn PFF treatment (400 nM, 24h) dampens presynaptic accumulation of VGLUT1 and VGAT induced by HA-NLG1A-B+ (**A**). In contrast, α-syn PFF treatment does not appear to affect HA-NLGN2-induced accumulation of VGLUT1 or VGAT (**D**). Treatment with α-syn monomers has no significant effects on VGLUT1 or VGAT accumulation induced by HA-NLG1A-B+ or HA-NLG2A+. Scale bars: 20 μm. (**B, C, E, F**) Quantification of presynaptic accumulation of VGLUT1 (**B, E**) and VGAT (**C, F**) in hippocampal neurons co-cultured with HEK293T cells expressing HA-NLG1A-B+ (**B, C**, HA-NLG2A+ (**E, F**), or HA-CD4, a negative control. Kruskal-Wallis one-way ANOVA, P < 0.0001. #P < 0.0001 compared with HA-CD4 and ****P < 0.0001 for the indicated comparisons with PBS control by Dunn’s multiple comparisons test. N.S., not significant. Data are presented as mean ± SEM. (n > 60 cells for each from four independent experiments).

## Discussion

In this study, we isolated β-NRXs as synaptic organizers that bind to α-syn PFFs, but not α-syn monomers, at the nanomolar range. We further determined that the HRD of β-NRXs is a domain responsible for α-syn PFF binding. Our gain-of-function assays using primary hippocampal neurons show that β-NRXs are involved in axonal binding of α-syn PFFs and that α-syn PFFs diminish β-NRX-mediated presynaptic organization. Thus, the binding of α-syn PFF to β-NRXs may impair synapse organizing activity of β-NRX-based synaptic organizing complexes. Given that β-NRXs play pivotal roles in synaptic transmission and plasticity (43), our results suggest that β-NRXs may act as a functional receptor for α-syn PFFs to mediate α-syn-induced synaptic pathology.

Although α-syn PFF pathology is proposed to be initiated from axons (14,57–59), little is known about presynaptic molecules that bind to α-syn PFFs. Previous studies have identified several α-syn PFF-binding membrane proteins such as PrP^c^, LAG-3 and FcγRIIB (47–49,60–62). PrP^c^ is localized to postsynaptic densities (63), but the subcellular localization of LAG-3 and FcγRIIB in neurons is undetermined. However, β-NRXs localize to presynaptic sites to function as synapse organizers (17,21,22,24,50). Our results demonstrating that β-NRXs bind to α-syn PFFs, but not α-syn monomers, that axonal overexpression of β-NRXs enhances the binding of α-syn PFFs on the axon surface, and that α-syn PFFs inhibit β-NRX-mediated presynaptic organization (inhibition of the synaptogenic activity of NLG1A-B+, which binds to only β-NRXs (55)) provide some of the first insights into presynaptic mechanisms of α-syn PFF pathology.

To further understand the molecular mechanism of the pathological pathway, we first investigated the specifics of β-NRX-α-syn PFF binding. β-NRXs possess an N-terminal domain called the histidine-rich domain (HRD) that is not present in α-NRXs (50). We have previously revealed that the HRD of β-NRXs is responsible for the binding of AβOs to β-NRXs (28,29). Our present study demonstrates that the HRD of β-NRXs is also responsible for the binding of α-syn PFFs. Thus β-NRXs bind both of the toxic protein aggregates central to two major types of neurogenerative disease: AβOs and α-syn PFFs in AD and PD, respectively (64). Interestingly, PrP^c^ also binds to both AβOs (63,65) and α-syn PFFs (47–49). The region of PrP^c^ responsible for binding to AβOs is the charge cluster region, an unstructured central domain containing many positively charged amino acid residues such as lysine (65). The HRD of β-NRXs is also an unstructured region containing clusters of positively charged amino acid residues including histidine and lysine (50,66). However, there is very low homology of the amino acid sequences of the PrP^c^ charge cluster region and the HRD of β-NRXs, suggesting that the unique electrical properties of the β-NRX HRD and the PrP^c^ central domain may be involved in the binding of abnormal protein aggregates related to neurodegenerative disorders. Further characterization of their electrical properties would be an interesting future study to better understand the molecular logic of their interactions and for therapeutics development.

We next determined the consequences of NRX-α-syn PFF binding on presynaptic organization induced by trans-synaptic organizing complexes. Our artificial synapse formation assays show that α-syn PFFs inhibit the accumulation of VGLUT1 and VGAT induced by NLG1A-B+. In contrast, α-syn PFFs affect neither VGLUT1 nor VGAT accumulation induced by NLG2A+. Given that NLG1A-B+ interacts with β-NRX but not α-NRXs whereas NLG2A+ interacts with both α- and β-NRXs because the splicing site B insert in NLGs prohibits the binding of α-NRXs (55) and NLG2 always lacks the splicing site B insert (17,22,50), these data suggest that α-syn PFFs selectively diminish β-NRX-mediated presynaptic organization. This raises the intriguing possibility that α-NRXs could compensate for α-syn PFF-induced diminished presynaptic organization, and future studies could investigate whether α-NRX-based interventions could counterbalance β-NRX-mediated α-syn PFF-induced synaptic toxicity.

Our previous study has shown that AβOs also diminish NRX-mediated presynaptic organization (28,29). However, the effects of AβOs on presynaptic organization are different from those of α-syn PFFs. One of the differences is that AβOs inhibit presynaptic organization induced by NLG2 (28), but α-syn PFFs do not. AβOs bind to not only β-NRXs but also α-NRXs that contain the splicing site 4 (α-NRX S4(+)) (28). Therefore, the binding of AβOs to α-NRXs may be involved in the inhibition of NLG2-induced presynaptic organization. Another major difference is that α-syn PFFs inhibit NLG1-induced accumulation of both VGLUT1 and VGAT whereas AβOs inhibit NLG1-induced accumulation of only VGLUT1, but not VGAT. Our cell surface protein binding assays show that α-syn PFFs bind to only β-NRXs, but AβOs bind to both β-NRXs and α-NRX S4(+). Thus, the NRX binding properties of α-syn PFFs and AβOs are not sufficient to explain the broader phenotype of α-syn PFFs induced effects on excitatory and inhibitory presynaptic organization. Another study has shown that the intracellular region of NRXs is dispensable to mediate NLG1-induced presynaptic organization (67), suggesting that other presynaptic transmembrane molecules that interact with the NRX ectodomain in cis may be involved in induction of presynaptic organization by NLG1 via the NRX ectodomain. Indeed, several transmembrane proteins that interact with the NRX ectodomain in cis have been isolated such as PTPσ (68,69), which interacts with NRXs via heparan sulfate chains on the NRX ectodomain, and SorCS1/2, which interacts with β-NRXs via the β-NRX HRD (70,71). Therefore, α-syn PFFs and AβOs might differentially affect cis-interactions between NRXs and NRX-interacting molecules, and this could be a basis for the synapsetype-specific pathological phenotypes of α-syn PFFs and AβOs.

Another remaining question is how α-syn PFFs diminish NLG1A-B+-induced presynaptic organization. One possibility is that α-syn PFFs might enhance the internalization, cleavage and/or degradation of β-NRXs and/or inhibit axonal transport of β-NRX. This would lead to reduction of axonal surface expression of β-NRXs, as occurs upon AβO exposure (28). However, given the distinct effects of AβOs and α-syn PFFs on VGLUT1 and VGAT accumulation, other underlying mechanisms should be considered for α-syn PFFs. For example, α-syn PFFs might block trans-interaction between β-NRXs and NLG1A-B+. Future studies are important to address what molecular and cellular mechanisms underlie the α-syn PFF inhibition of NLG1-induced presynaptic organization.

A major physiological role of β-NRXs in hippocampal neurons is to control glutamate release probability through endocannabinoid signaling pathways (43). A previous study has shown that in cultured hippocampal neurons, α-syn PFF treatment decreases the frequency of miniature excitatory postsynaptic currents (mEPSC), but not mEPSC amplitude (72), suggesting that α-syn PFFs downregulate glutamate release probability. Further, dopaminergic neurons and striatal neurons are also regulated by endocannabinoid signaling pathways (73). Indeed, pharmacological manipulations of endocannabinoid signaling pathways have attracted attention as potentially therapeutic for PD and have been tested in clinical trials (74,75). It would therefore be interesting to study whether and how α-syn PFFs affect endocannabinoid signaling pathways through β-NRXs to induce synaptic dysfunction and/or neuronal toxicity in synucleinopathies.

Our present study suggests that shielding β-NRXs from α-syn PFF binding would be helpful for alleviating α-syn PFF pathology. We have recently revealed that the protein sorting receptor SorCS1 shields β-NRXs from AβO binding and rescues AβO-induced synaptic pathology (71). Of note, SorCS1 also binds to β-NRXs through the β-NRX HRD to compete with AβOs for β-NRX interaction (71). Given that like AβOs and SorCS1, α-syn PFFs bind to β-NRXs through the HRD, it is possible that SorCS1 could also shield β-NRXs from α-syn PFF binding to rescue α-syn PFF-induced synaptic and neuronal toxicity. Another strategy to shield β-NRXs from α-syn PFF binding would be using the anti-NRX1β HRD antibody that we showed effectively inhibits α-syn PFF binding to NRX1β in this study. Thus, in future studies, it will be important to address whether and how SorCS1 overexpression and anti-NRX1β treatment rescue α-syn PFF-induced pathology in culture and *in vivo* for the development of novel therapeutic strategies for synucleinopathies.

## Experimental procedures

### Plasmids

The following deletion constructs and point mutations for extracellular HA-tagged NRX1βS4(-) (NP_001333889.1) (HA-NRX1βS4(-)) were made by inverse PCR with the vector expressing HA-NRX1βS4(-) under CMV promoter as a PCR template followed by DpnI digestion: ΔLNS (amino acids (aa) 107-236 deleted), ΔCysloop (aa 319-329 deleted), △HS (S316A). For the internal ribosome entry site (IRES)-based bicistronic constructs for co-expressing GFP with either HA-NRX1βS4(-) or HA-NRX1βΔHRDS4(-) under the CAG promoter (pCAG-HA-NRX1βS4(-)-IRES-GFP and pCAG-HA-NRX1βΔHRDS4(-)-IRES-GFP, respectively), we amplified the coding sequences of HA-NRX1βS4(-) and HA-NRX1βΔHRDS4(-) by PCR using pCMV-HA-NRX1βS4(-) and pCMV-HA-NRX1βΔHRDS4(-) that we generated before (28) as templates and then subcloned the PCR products into the pCAG-IRES-GFP vector (71) at the EcoRI site. The following plasmids were kind gifts: pCAG-HA-NRX1βS4(-) from Dr. Takeshi Uemura (Shinshu University); HA-NLGN1A(-)B(-), HA-NLGN1A(+)B(-), HA-NLGN1A(-)B(+), HA-NLGN1A(+)B(+), HA-NLG3 and HA-NLG4 from Dr. Peter Scheiffele (University of Basel) via Addgene; HA-NLGN2, from Dr. Ann Marie Craig (University of British Columbia); HA-glutamate receptor delta-1 (GluD1) and HA-GluD2 from Dr. Michisuke Yuzaki (Keio University). The other constructs used in this study were described previously (28,71,76–78). All constructs were verified by DNA sequencing.

### Generation of α-synuclein PFFs and biotin labelling

Using conventional methods for bacterial transformation and large-scale protein expression, BL21 (DE3) Escherichia coli were transformed with pGEX-6-alpha synuclein, a plasmid containing a sequence encoding glutathione S-transferase (GST)–tagged full-length recombinant human α-synuclein (NM_000345) and with a plasmid containing GST-tagged recombinant 3C enzyme, which can cleave the GST tag from GST-tagged proteins. pGEX-6-alpha synuclein (backbone plasmid: pGEX6P1) was originally purchased from the University of Dundee MRC Protein Phosphorylation and Ubiquitination Unit (#DU30005). The plasmid used to express GST-3C protease was created by cloning the coding sequence for human rhinovirus (HRV) 3C protease into the pGEX-2T backbone plasmid. The GST-tagged proteins were purified from the bacterial cell lysates by affinity column chromatography. The purified GST-α-synuclein protein was then treated with the purified GST-3C enzyme to remove the GST tag. Untagged α-synuclein protein was purified from the reaction using a GSTrap 4B column and then further purified using size exclusion chromatography with a Superdex 200 16/600 column on the ÄKTA pure L system. Afterwards, human α-syn fibrils were generated from aliquots (500 μL of 5 mg/ml in PBS) of recombinant α-syn monomers that were shaken at 1000 rpm in a ThermoMixer at 37 °C for 5 days. The generated fibrils were sonicated for 30s at 10% power with a 50% duty cycle (0.5s on, 0.5s off). The resulting preformed fibrils (PFFs) were labeled with biotin by using an EZ-Link™ Sulfo-NHS-LC-Biotinylation Kit (Cat#21435, Thermofisher Scientific). After biotinylation, the PFFs were purified using Zeba™ Spin Desalting Columns (7K MWCO, Cat#89882, Thermofisher Scientific) to remove excess of unbound biotin and stored at −80 °C.

### Validation of α-synuclein PFFs formation by electron microscopy, dynamic light scattering and thioflavin-T assays

Electron microscopy (EM), dynamic light scattering (DLS) and thioflavin-T (ThT) analyses were used as quality controls for α-syn PFFs validation. Briefly, EM, samples containing 20 μM (monomer equivalent) α-syn aggregates before and after the sonication step (see the above *Generation of α-synuclein preformed fibrils* section) were prepared by dilution in ddH_2_O. 5 μl of each preparation was deposited on copper-coated grids for 2 mins. Then, 4% PFA was added onto the grids for 1 min and then the grids were washed three times using ddH_2_O. Finally, 2% uranyl acetate was added onto the grids before covering them with a glass dish. The grids were then visualized under an electron microscope. Analysis of the images was performed using Fiji-ImageJ and MatLab R2017b software. For DLS analysis, a ≥0,6 mg/ml α-syn PFFs solution was prepared by dilution in PBS. After a centrifugation step (13000 RPM, 5 mins), the supernatant was transferred to a new tube and analyzed using the Zetasizer Nano S system. For ThT assays, 300 μl of PFF sample (~50 μg/ml) was mixed with 300 μl of 25 μM ThT solution at room temperature for 20 min, then 100-μl aliquots were transferred to a Corning 96-well plate (black with clear bottom), and fluorescent signals were read by a microplate reader with excitation at 450 nm and emission at 490 nm, as previously described (45), with PBS (vehicle) and monomeric α-syn as controls.

### Dot blot

For dot blot experiments, 1 μg of either α-syn PFFs or monomers (biotinylated or untagged) were applied in spots on a nitrocellulose membrane (Cat#1620145, BioRad) for 1h at room temperature (RT). The membrane was exposed to a TBST/BSA 5% blocking solution (8 g NaCl, 200 mg KCl, 3 g Tris HCl, 0,05% Tween20 and 5% m/v BSA) for 30 mins at RT under gentle agitation then a primary antibody against α-syn diluted in TBST/BSA 0,1% for 2 h at RT, followed by washing and finally treatment with an HRP-conjugated secondary antibody. The membrane was then exposed to ECL detection reagent (Cat#1705061, BioRad) for 5 mins at RT and detection was performed using a Chemidoc XRS+ system (BioRad). After a 30-min stripping step (1M Glycine HCl pH 2,7, 20% SDS in MiliQ), the membrane was exposed to HRP-conjugated streptavidin for 1h at RT followed by ECL detection reagent application and image acquisition. The specific reagents used were a primary antibody directed against α-syn (1:2000; mouse IgG, Cat#32-8100, Thermofisher Scientific), an HRP-conjugated anti-mouse secondary antibody (1:5000; donkey host, Cat#715-035-151, Jackson ImmunoResearch), and HRP-conjugated streptavidin (1:10000; Cat#21130, Pierce™ High Sensitivity Streptavidin-HRP, Thermofisher Scientific).

### Cell surface binding assays

To test for interaction of biotin-α-syn PFFs or biotin-α-syn monomers with our proteins of interest, COS-7 cells cultured on coverslips were transfected with the indicated expression vectors using TransIT-LT1 Transfection Reagent (Mirus bio) and maintained for 24 h. The transfected COS-7 cells were washed with extracellular solution (ECS) containing 2.4 mM KCl, 2 mM CaCl_2_, 1.3 mM MgCl_2_, 168 mM NaCl, 20 mM HEPES (pH 7.4) and 10 mM D-glucose supplemented with 100 μg/ml of bovine serum albumin (BSA; ECS/BSA). Next, the transfected COS-7 cells were incubated with biotin-α-syn PFFs or monomers previously diluted at the indicated concentration in ECS/BSA and were kept for 1h at 4 °C to prevent endocytosis. For the competition experiment, anti-NRX1β antibody (mouse IgG, N170A/1, NeuroMab) recognizing the HRD of the NRX1β was added to the ECS/BSA solution containing biotin-α-syn at the indicated concentrations. The cells were washed three times using ECS/BSA and two times using ECS solution, then fixed using parafix solution (4% paraformaldehyde and 4% sucrose in PBS [pH 7.4]) for 12 min at RT. To label surface HA or bound biotin-α-syn, the fixed cells were then incubated with blocking solution (PBS + 3% BSA and 5% normal donkey serum) for 1 h at RT. Afterwards, they were incubated with primary antibodies in blocking solution overnight at 4 °C and with fluorescent-conjugated secondary antibodies and/or fluorescent-conjugated streptavidin to label bound biotin-α-syn for 1 h at RT. The specific reagents used were an anti-HA primary antibody (1:2000; rabbit IgG, Cat# ab9110, Abcam), a highly cross-absorbed Alexa488-conjugated donkey anti-rabbit IgG (H+L) secondary antibody (1:500; Jackson ImmunoResearch), and Alexa594-conjugated streptavidin (1:2500; Jackson ImmunoResearch).

### Surface plasmon resonance

SPR analyses were performed using a Biacore T200 instrument (GE Healthcare). Purified recombinant NRX1β ectodomain fused to human immunogloblin Fc (NRX1β-Fc, R&D sytems, Cat#5268-NX-05, R&D system) was immobilized on carboxymethylated dextran CM5 sensor chips (Cytiva) using an amine-coupling strategy. Briefly, the sensor chip surface was activated with a 1:1 mixture of N-hydroxysuccinimide and 3-(N,N-dimethylamino)-propyl-N-ethylcarbodiimide. NRX1β-Fc solution (solubilized in acetate buffer, pH 5.0) was injected at a flow rate of 20 μl/min in HBS-N running buffer (10 mM HEPES, 150 mM NaCl, pH 7.4, 0.05% [v/v] Tween-20) to reach a level of immobilization of 300 relative units (RU) on the CM5 sensor chip. Surfaces (protein and reference) were blocked by the injection of an ethanolamine solution. The binding kinetics of α-syn PFFs and α-syn monomers over the NRX1 β sensor chip were evaluated in HBS-N running buffer (10 mM HEPES, 150 mM NaCl, pH 7.4, 0.05% [v/v] Tween-20), with concentrations ranging from 0 to 10 μM. All tests were performed at 25 °C using a flow rate of 30 μl/min. Sensor chip surfaces were regenerated by injecting 15 μl of a 10 mM Glycine pH 3.0 solution at a flow rate of 30 μl/min. Binding sensorgrams were obtained by subtracting the reference flow cell data. Data analysis was performed using BIA Evaluation Software (GE Healthcare) and fit to a one-site Langmuir adsorption model.

#### Neuron culture, transfection, and neuronal immunocytochemistry

Primary rat hippocampal neuron cultures were prepared from embryonic day 18 (E18) rat embryos as described previously (79). All animal experiments were carried out in accordance with the Canadian Council on Animal Care guidelines and approved by the Institut de Recherches Cliniques de Montréal (IRCM) Animal Care Committee. Dissociated hippocampal cells were plated onto coverslips in two different manners: in wells of 12-well plates containing coverslips at a density of 200,000 neurons/coverslip (referred to as high-density neuron cultures) or on 6-cm dishes each containing five coverslips at a density of 300,000 neurons/dish (referred to as low-density neuron cultures). 24h after plating, coverslips containing low-density neurons were transferred to another 6-cm dish containing a glial feeder layer. To verify the pathogenicity of the α-syn PFFs preparation, low-density hippocampal neurons were treated at 8 days *in vitro* (DIV) with α-syn PFFs or α-syn monomers (2.5 μg/ml) for 7 days. For the cell surface binding assays in low-density transfected neurons, the transfection was performed at 9 DIV using the ProFection® Mammalian Transfection System from Promega (Cat#E1200). At 15 DIV, neurons were treated with biotin-α-syn PFFs (2.5 μg/ml) for 1h at 37 °C. At the end of the experiments, neurons were fixed with parafix solution for 12 min, permeabilized with PBST (1xPBS + 0.2% Tween20) [except for experiments examining surface expression], and then blocked with blocking solution. Afterwards, they were incubated with primary antibodies in blocking solution overnight at 4 °C and with secondary antibodies for 1 h at RT. To label surface HA-tagged constructs and biotin-α-syn, together with MAP2, the fixed neurons were incubated sequentially with a primary antibody against HA and dye-conjugated streptavidin without cell permeabilization and then permeabilized with PBST for MAP2 immunostaining. The following primary antibodies were used for immunocytochemistry: anti-HA (1:2000; rabbit IgG, Cat#ab9110, Abcam), anti-MAP2 (1:2000; chicken polyclonal IgY; Cat#ab5392, Abcam), anti-HA (1:2000; mouse IgG2bκ, Cat#11583816001, Millipore Sigma), anti-VGLUT1 (1:250; guinea pig; Cat#135304, Synaptic System), anti-VGAT (1:1000; Rabbit IgG, Cat#131 003, Synaptic System) and anti-α-syn [phospho S129] (1:1000; mouse IgG, Cat#ab184674, Abcam). Highly cross-adsorbed Alexa dye-conjugated secondary antibodies generated in donkey toward the appropriate species (1:500; Alexa488, Alexa594, and Alexa647; Jackson ImmunoResearch) were used as detection antibodies. For biotin detection, Alexa594-conjugated streptavidin (1:2500) was used.

### Artificial synapse formation assays

The artificial synapse formation assays were performed as we did previously (28,78). Briefly, HEK293T cells were first transfected as described in the cell surface binding assay section. After 24h, the cells were harvested through trypsinization, and 10,000 cells were added to 15-DIV high-density neuron cultures simultaneously with α-syn PFFs (400 nM monomer equivalent), monomers (400 nM) or PBS. After a 24-hour incubation period, neurons were fixed using parafix solution for 12 mins at RT and blocked using a blocking solution. To identify surface HA-tagged protein expression, cells were treated with a primary antibody against HA overnight at 4 °C. Next, neurons were treated with PBST, blocking solution, and stained for MAP2, VGLUT1 and VGAT overnight at 4 °C using corresponding primary antibodies.

### Imaging and quantitative fluorescence analysis

For quantitative analysis, all image acquisition was performed on a Leica DM6000 fluorescent microscope with a 40 × 0.75 numerical aperture (NA) dry objective or 63 × 1.4 NA oil objective and a Hamamatsu cooled CCD camera using Volocity software (Perkin Elmer). Images were obtained in 12-bit grayscale and prepared for presentation using Adobe Photoshop 2020. The only exception is the cell surface binding assays on neurons for which images were captured using a Leica SP8 confocal microscope with a 63x objective. For quantification, sets of cells were immunostained simultaneously and imaged with identical microscope settings. Analysis for the COS-7 based cell surface binding assays was performed using Volocity, and that for the other assays was performed using Metamorph 7.8 (Molecular Devices). For the cell surface binding assays, after off-cell background intensity was subtracted, the average intensity of bound proteins per COS-7 cell region was measured and normalized to the average intensity of the surface HA signal. The half-maximal inhibitory concentration (IC_50_ value) was determined by non-linear regression curve fitting in GraphPad Prism9 (GraphPad Software). For the artificial synapse formation assays, images were acquired where HA-expressing HEK293T cells were observed and the VGLUT1 and VGAT total intensity around the HEK293T cells was calculated after thresholding the background. For cell surface binding assays on neurons, GFP-positive but MAP2-nagative neurites were selected as axons. Using Metamorph 7.8 (Molecular Devices), the axons were traced by a line command, and the average intensity value of bound α-syn PFFs and that of surface HA along the selected line regions were measured by a region measurement command.

### Statistical analysis

Statistical tests were performed using GraphPad Prism 9 (GraphPad Software). The majority of the data had non-normal distribution and non-equal variance, so statistical comparisons were made using Kruskal-Wallis one-way analysis of variance (ANOVA) with post-hoc Dunn’s multiple comparisons test. Data are reported as the mean ± the standard error of the mean (SEM) from three independent experiments, if not stated otherwise. Statistical significance was defined as P < 0.05.

## Data Availability

This study includes no data deposited in external repositories.

## Acknowledgments

We thank Emmanuelle Nguyen-Renoufor for her technical assistance. This study was supported by a Parkinson Canada New Investigator Award (2018-00079), a Canadian Institutes of Health Research (CIHR) Project Grant (PTJ-159588) and Fonds de la recherche du Québec–Santé (FRQS) Research Scholars (Junior 2-29106 and Senior-251655) to H.T, as well as by funding from the Canada First Research Excellence Fund, awarded through the Healthy Brains, Healthy Lives initiative at McGill University to T.M.D, a Natural Sciences and Engineering Research Council of Canada (NSERC) Discovery Grant (RGPIN-2018-06209) to S.B., an FRQS and Parkinson Québec Master’s training award (273951), a CIHR Canada Graduate Scholarships – Master’s Award and Université de Montréal Bourse de recrutement du Département de neurosciences to A.F., an NSERC doctoral award, an FRQS doctoral award (252652), an IRCM doctoral scholarship and a McGill medical internal studentship to A.K.L.

## Authors’ contributions

B.F. performed a majority of the experiments. A.F. and A.L. initiated this project by performing pilot experiments. W.L, I.S. and T.M.D. worked on the generation and validation of α-synuclein proteins. P.T.N and S.B. worked on the SPR experiments. S.K. performed some cell surface binding assays. H.T. conceived and supervised the project. B.F. and H.T. prepared the manuscript with contributions from the other authors.

