## Supplementary Figure 1, 2 and 3 for "α-synuclein preformed fibrils bind to β-neurexins and impair β-neurexin-mediated presynaptic organization"

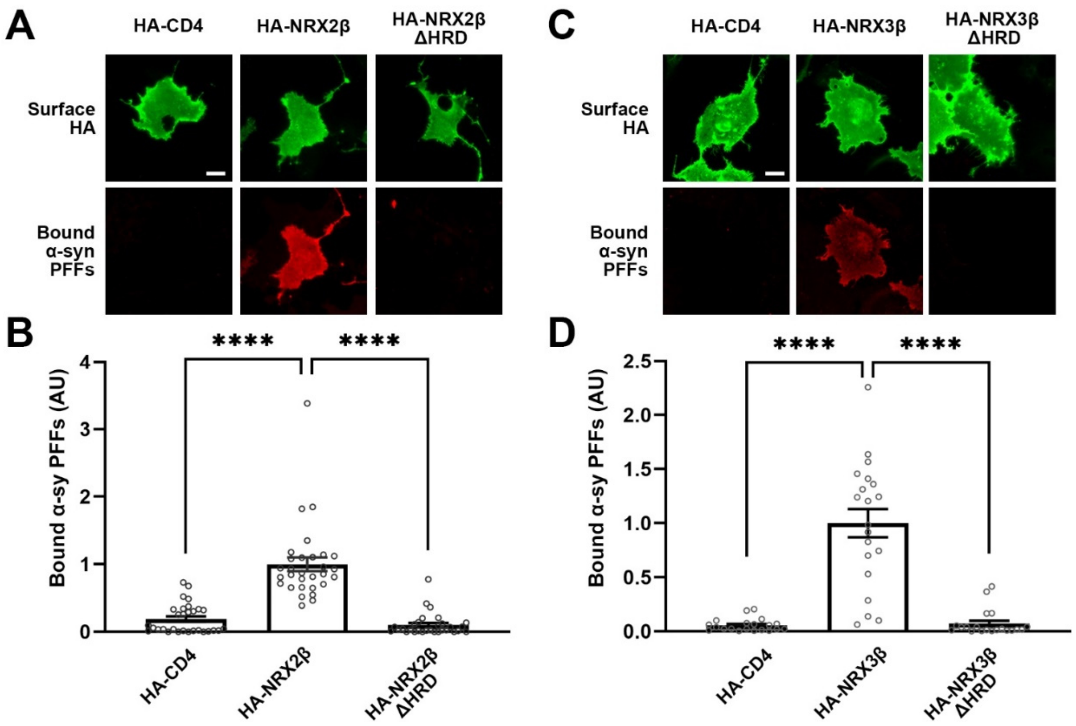


**Supplementary Figure 1. The N-terminal histidine-rich domain of neurexin2β and 3β is responsible for the binding of α-syn PFFs.**

(**A, C**) Representative images showing the binding of 1 μM biotin-α-syn PFFs to COS-7 cells expressing the indicated HA-NRX2β (**A**) and HA-NRX3β (**C**) constructs, either full-length or lacking the histidine-rich domain (HRD) (ΔHRD), or HA-CD4, a negative control. COS-7 cells expressing HA-NRX2β ΔHRD or HA-NRX3β ΔHRD show no binding of biotin-α-syn PFFs, appearing comparable to those expressing HA-CD4. All constructs have significant surface HA expression.

(**B, D**) Quantification of the average intensity of bound biotin-α-syn PFFs on COS-7 cells expressing the indicated HA-NRX2β/3β constructs or HA-CD4. Kruskal-Wallis one-way ANOVA, $P < 0.0001$. ****$P < 0.0001$ in the indicated comparisons by Dunn's multiple comparisons test. N.S., not significant. Data are presented as mean ± SEM. ($n = 30$ and 20 cells for **B and D** from three and two independent experiments, respectively).

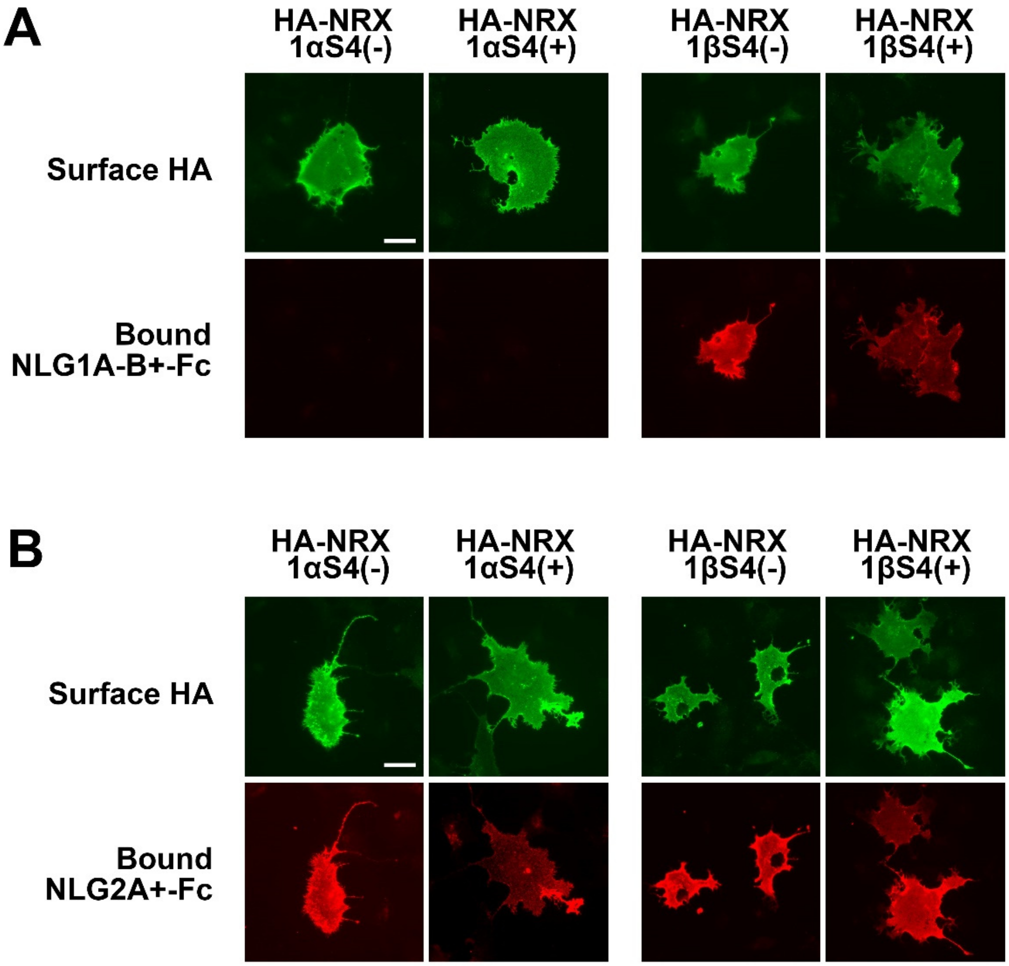


**Supplementary Figure 2. NLG1A-B+ binds to NRX1β but not NRX1α whereas NLG2A+ binds to both NRX1α and NRX1β.**

(**A, B**) Representative images showing the binding of NLG1A-B+-Fc (**A**) and NLG2A+-Fc (**B**) to COS-7 cells expressing the indicated HA-NRXs constructs. As reported previously (22,55), NLG1A-B+-Fc binds to only NRX1β, and not NRX1α. In contrast, NLG2A+-Fc binds to both NRX1α and 1β. Scale bars: 30 μm.

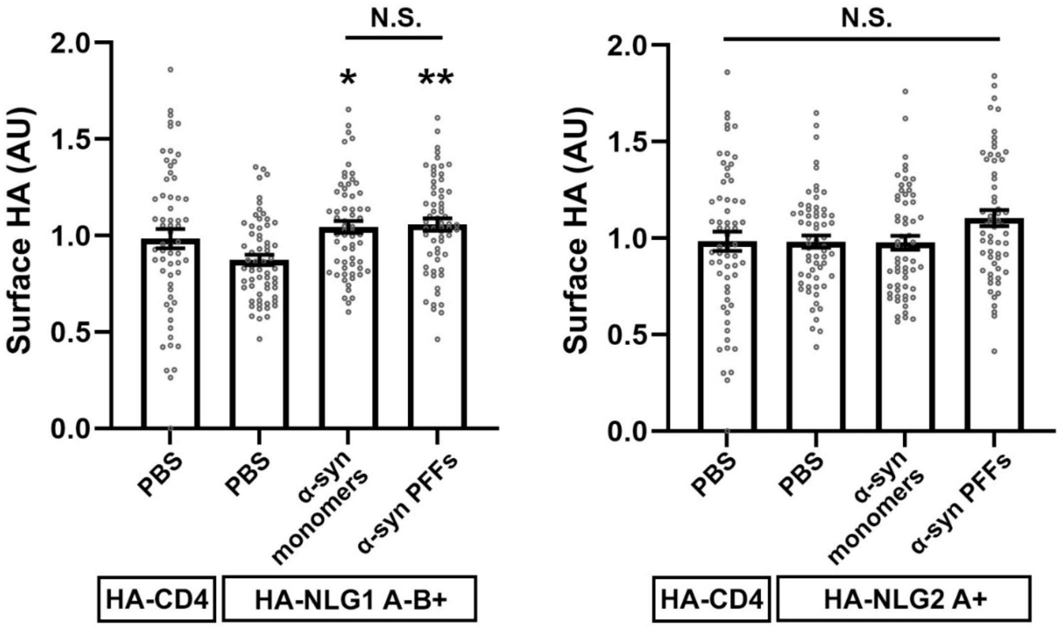


**Supplementary Figure 3. α-syn PFF treatment does not decrease surface expression of HA-NLG1A-B+ or HA-NLG2A+ on HEK293T cells in the co-culture-based artificial synapse formation assays presented in Figure 7.**

Quantification of surface HA expression level of HA-NLG1A-B+ (left) and HA-NLG2A+ (right) under the indicated treatment conditions in the co-culture-based artificial synapse formation assays presented in Figure 7. The level of surface HA expression after α-syn PFF treatment in the NLG1A-B+ group (left) does not correlate with the NLG1A-B+-induced VGLUT1 and VGAT accumulation phenotype after α-syn PFF treatment (**Fig. 7B and E**), excluding the possibility that the phenotype is due to decreased surface expression of HA-NLG1A-B+ on HEK293T cells after α-syn PFF treatment. Kruskal-Wallis one-way ANOVA, $P < 0.001$ for HA-NLG1 and $P = 0.1439$ for HA-NLG2A+. *$P < 0.01$ and **$P < 0.001$ for the indicated comparisons by Dunn's multiple comparisons test. N.S., not significant. Data are presented as mean ± SEM. ($n > 60$ cells for each from four independent experiments).
